# AlphaMissense is better correlated with functional assays of missense impact than earlier prediction algorithms

**DOI:** 10.1101/2023.10.24.562294

**Authors:** Alicia Ljungdahl, Sayeh Kohani, Nicholas F. Page, Eloise S. Wells, Emilie M. Wigdor, Shan Dong, Stephan J. Sanders

## Abstract

Missense variants that alter a single amino acid in the encoded protein contribute to many human disorders but pose a substantial challenge in interpretation. Though these variants can be reliably identified through sequencing, distinguishing the clinically significant ones remains difficult, such that “Variants of Unknown Significance” outnumber those classified as “Pathogenic” or “Likely Pathogenic.” Numerous *in silico* approaches have been developed to predict the functional impact of missense variants to inform clinical interpretation, the latest being AlphaMissense, which uses artificial intelligence methods trained on predicted protein structure. To independently assess the performance of AlphaMissense and 38 other predictors of missense severity, we compared predictions to data from multiplexed assays of variant effect (MAVE). MAVE experiments generate almost every possible individual amino acid change in a gene and measure their functional impact using a high-throughput assay. Assessing 17,696 variants across five genes (*DDX3X, MSH2, PTEN, KCNQ4*, and *BRCA1*), we find that AlphaMissense is consistently one of the top five algorithms based on correlation with functional impact and is the best-correlated algorithm for two genes. We conclude that AlphaMissense represents the current best-in-class predictor by this metric; however, the improvement over other algorithms is modest. We note that multiple missense predictors, including AlphaMissense, appear to overcall variants as pathogenic despite minimal functional impact and that substantially more high-quality training data, including consistently analyzed patient cohorts and MAVE analyses, are required to improve accuracy.

## Introduction

Missense variants alter single amino acids in protein-coding genes and are a major cause of human morbidity and mortality. With advances in sequencing, almost all of the ∼8,500 missense variants in each individual can be identified reliably. However, distinguishing the one or two that might have a substantial clinically significant impact is a needle-in-a-haystack problem. Clinical variant interpretation uses the ACMG criteria to distinguish the “Pathogenic” from “Benign,” but thousands of individuals receive a “Variant of Unknown Significance” report instead.^1^ The challenge of variant classification already hinders diagnosis, prognosis, management, and family counseling; as genome-targeted therapies develop, this interpretive uncertainty will become even more acute.

For proteins with well-characterized biological actions, functional assays can be developed to assess the impact of individual missense variants. This process is often laborious and must be adapted to specific genes due to substantial diversity in proteomic function. Even in the era of genomics, the vast majority of missense variants remain uncharacterized. Of those that have been characterized, it is apparent that most missense variants in most genes have minimal impact. However, there remain many missense variants that lead to varying degrees of loss-of-function, and a small number lead to altered function (e.g., a gain-of-function or dominant negative effect). Further complicating this landscape, some proteins have multiple roles, so a missense variant’s impact could be specific to one or many of these roles; furthermore, some missense variants can cause mixed loss-of-function and gain-of-function effects.^2^

Given this background, a predictive computational approach to characterize missense variants is highly attractive. Early approaches, such as the Grantham Score,^3^ developed five decades ago, assessed biochemical differences in amino acids. Major advances have included the use of cross-species conservation data (e.g., PolyPhen^4^), integration of multiple scores to make ensemble predictors (e.g., REVEL^5^), leveraging patterns of genomic constraint in humans (e.g., MPC^6^), and applying artificial intelligence approaches (e.g., PrimateAI-3D^7^). Recently, AlphaMissense was released,^8^ bringing proteome-wide structural predictions to bear on this problem. The authors’ comparison to categorically classified ClinVar variants shows evidence of AlphaMissense outperforming other missense severity scores.

Here, we performed an orthogonal and independent assessment of AlphaMissense and 38 other missense severity predictors using continuously distributed functional data from previously published multiplexed assays of variant effect (MAVE) of 17,696 missense variants in five genes, *DDX3X*,^9^ *BRCA1*,^10^ *MSH2*,^11^ *PTEN*,^12^ and *KCNQ4*.^13^ These MAVE approaches use genomic technologies (e.g., saturation mutagenesis) and high-throughput or competitive assays to systematically assess thousands of missense variants in parallel. *DDX3X* is an X-linked RNA helicase associated with neurodevelopmental delay; missense function was assessed using saturation CRISPR-Cas9 homology-directed repair (HDR) in the HAP1 haploid human cell line, in which DDX3X is an essential gene, so reduced function impaired cell survival.^9^ Both *BRCA1* and *MSH2* are involved in DNA repair and are associated with an increased risk of cancer.^10,11^ *BRCA1* function was assessed via cell survival in a genome-editing HDR model that disrupted and then attempted to rescue *BRCA1* with each variant in HAP1.^10^ A similar approach was used to assess *MSH2*, with the ability of cDNA containing each variant to induce sensitivity to 6-thioguanine (a toxin only in cells with functional mismatch repair) in a HAP1 *MSH2* knockout; here non-functional variants resulted in impaired cell survival.^11^ *PTEN* is a key enzyme in the phosphoinositide signaling pathway, converting phosphatidylinositol (3,4,5)-triphosphate (PIP_3_) into phosphatidylinositol (4,5)-bisphosphate (PIP_2_). Deficits in this lipid phosphatase activity lead to neurodevelopmental delay, macrocephaly, and cancer predisposition.^12^ *PTEN* function was assessed in an artificial humanized yeast model in which PIP_3_ concentration inhibits replication, so decreased replication rate reflects an inability to convert PIP_3_ to PIP_2_.^12^ *KCNQ4* is a potassium channel associated with deafness. The impact of variants in *KCNQ4* on the encoded K_V_7.4 potassium channel was assayed using whole-cell currents measured by patch clamp in CHO-K1 cells.^13^

## Methods

For each gene, all possible missense variants from single nucleotide variants were predicted from the Matched Annotation from NCBI and EMBL-EBI (MANE) isoform.^14^ Functional data were downloaded from the published manuscripts and annotated against 36 predictors of missense severity using ANNOVAR (dbnsfp42a).^15^ In addition, missense severity scores were added from EVE,^16^ PrimateAI-3D,^7^ and AlphaMissense,^8^ along with pathogenicity, as defined by ClinVar (downloaded October 2023). Correlation between functional scores and each of the 39 predictors of missense severity was assessed using absolute Spearman correlation. For *KCNQ4*, a gene in which variants could increase or decrease protein function, the absolute deviation from typical values was calculated and log-scaled to estimate absolute Spearman correlation.

## Results

Across 39 algorithms (Table S1), AlphaMissense was the highest correlated algorithm to the functional data for two genes (*PTEN, BRCA1*), 2^nd^ for *MSH2*, 3^rd^ for *DDX3X*, and 5^th^ for *KCNQ4* (Table 1, Fig. 1). No other algorithm was ranked in the top five for all genes; however, PrimateAI-3D was ranked in the top five for four and ranked first for two genes (*MSH2, KCNQ4*). These results are in line with the comparisons performed in the AlphaMissense manuscript.^8^

**Table 1.**
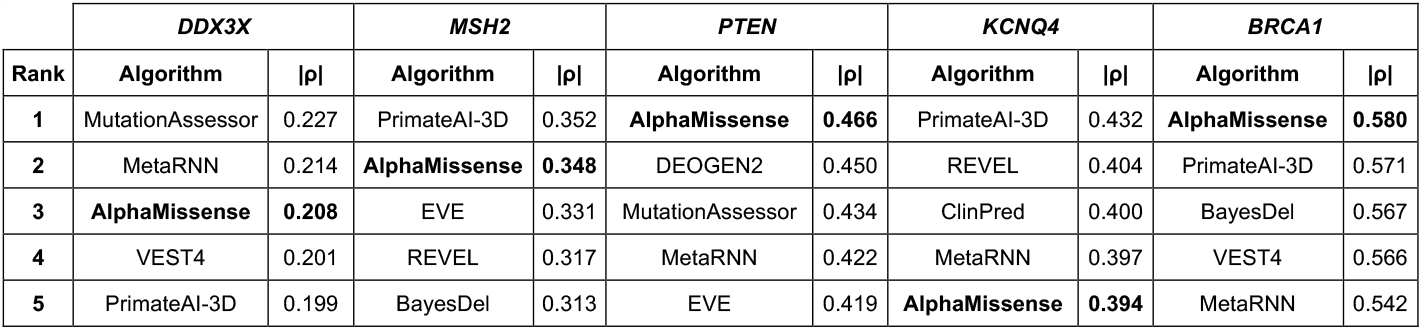
Top five out of 39 missense prediction algorithms by absolute Spearman correlation.

|  | *DDX3X* |  | *MSH2* |  | *PTEN* |  | *KCNQ4* |  | *BRCA1* |  |
| --- | --- | --- | --- | --- | --- | --- | --- | --- | --- | --- |
| Rank | Algorithm | $\vert \rho\vert $ | Algorithm | $\vert \rho\vert $ | Algorithm | $\vert \rho\vert $ | Algorithm | $\vert \rho\vert $ | Algorithm | $\vert \rho\vert $ |
| 1 | MutationAssessor | 0.227 | PrimateAI-3D | 0.352 | **AlphaMissense** | **0.466** | PrimateAI-3D | 0.432 | **AlphaMissense** | **0.580** |
| 2 | MetaRNN | 0.214 | **AlphaMissense** | **0.348** | DEOGEN2 | 0.450 | REVEL | 0.404 | PrimateAI-3D | 0.571 |
| 3 | **AlphaMissense** | **0.208** | EVE | 0.331 | MutationAssessor | 0.434 | ClinPred | 0.400 | BayesDel | 0.567 |
| 4 | VEST4 | 0.201 | REVEL | 0.317 | MetaRNN | 0.422 | MetaRNN | 0.397 | VEST4 | 0.566 |
| 5 | PrimateAI-3D | 0.199 | BayesDel | 0.313 | EVE | 0.419 | **AlphaMissense** | **0.394** | MetaRNN | 0.542 |

**Figure 1.**
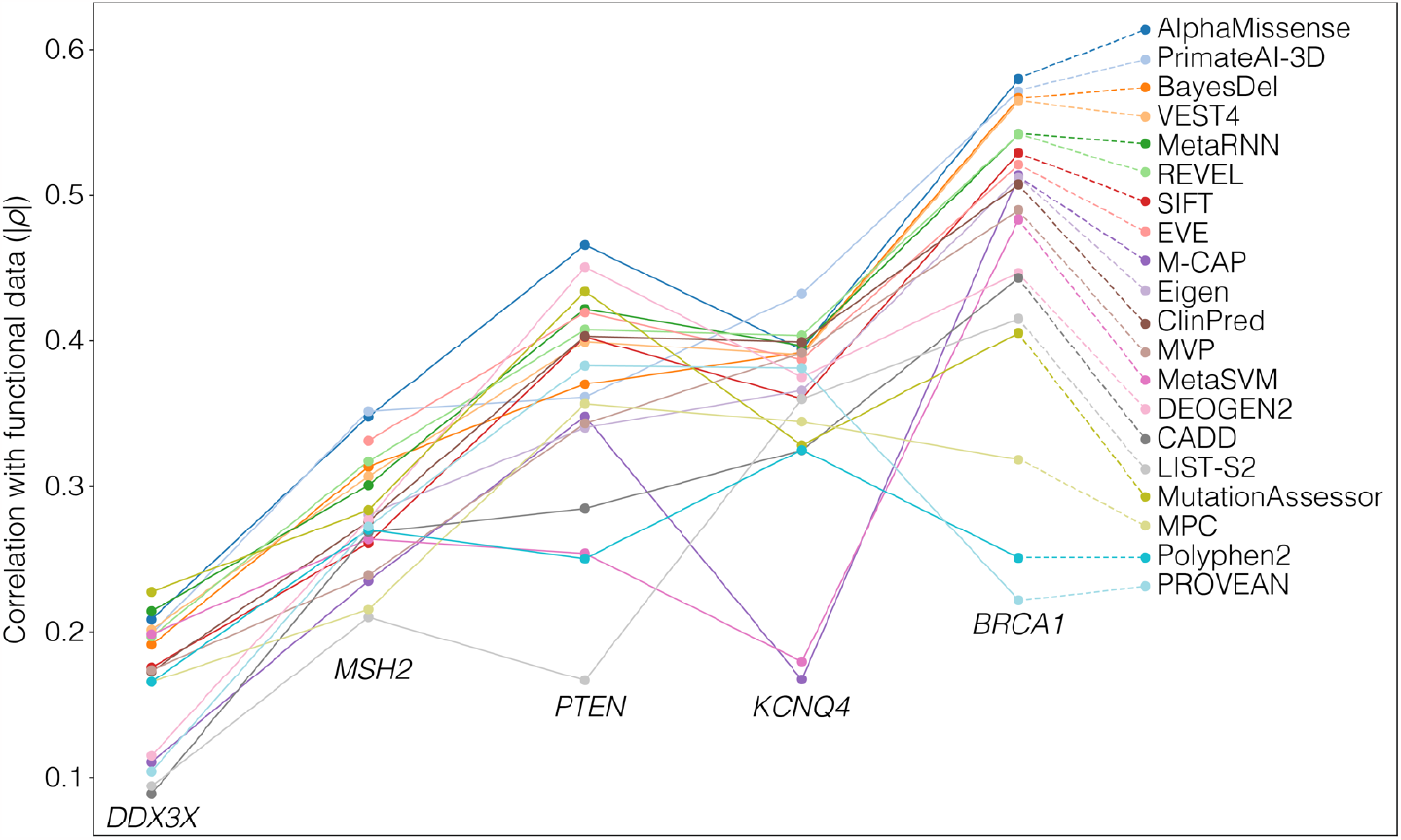
Comparison of functional data for 17,696 missense variants across 20 prediction algorithms. Data were collated for five genes: *DDX3X* (4,261 variants), *MSH2* (5,271 variants), *PTEN* (2,510 variants), *KCNQ4* (3,757 variants), and *BRCA1* (1,897 variants). The absolute Spearman correlation (|ρ|) between the functional data and the predicted scores was calculated in each of the five genes for 39 algorithms, of which 20 high-ranking and distinct algorithms are shown here (split by color).

To better understand the performance of AlphaMissense, we plotted the functional data against the predictions of both AlphaMissense and REVEL (an algorithm widely used for clinical predictions) for all variants in the five genes. In four of the five genes, AlphaMissense scores show a modestly higher correlation with the functional data than REVEL, while in *KCNQ4*, correlation is marginally higher with REVEL scores (Fig. 2). Both loss-of-function and gain-of-function variants in *KCNQ4* received high AlphaMissense and REVEL scores. Unsurprisingly, AlphaMissense and REVEL scores are highly correlated (Fig. 2D), and this was also true of AlphaMissense and other high-ranking algorithms (Table S1).

**Figure 2.**
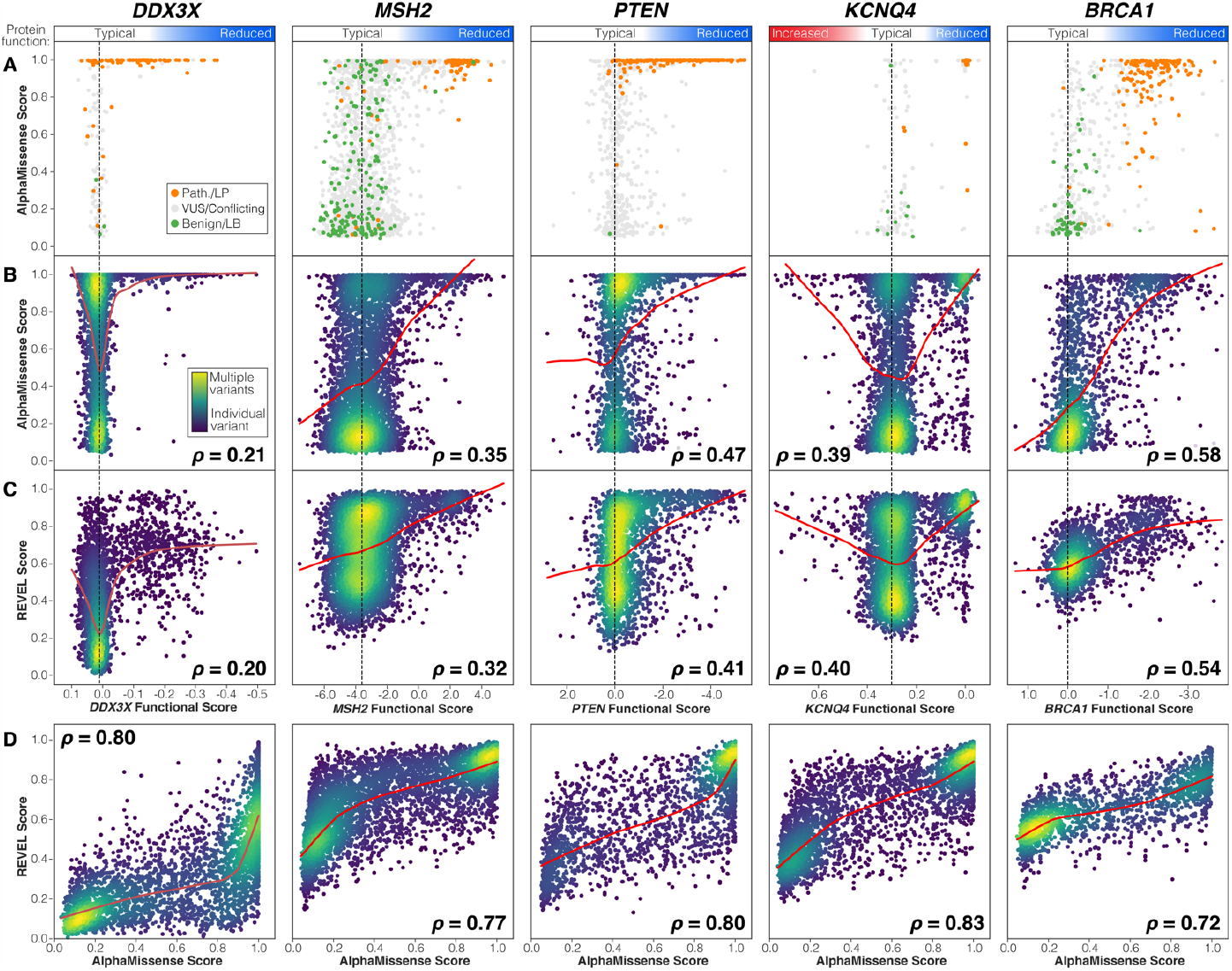
Comparison of functional data for 17,696 missense variants to ClinVar, AlphaMissense, and REVEL. **A)** For variants in ClinVar, functional data were compared to AlphaMissense score with points colored by ClinVar status (Pathogenic/Likely Pathogenic in orange, Variants of Unknown Significance and Conflicting Interpretations in grey, Benign/Likely Benign in green). **B)** For all variants, functional data were compared to AlphaMissense score. Point color represents variant density from high (yellow) to medium (green) to individual missense variants (dark blue). The locally weighted scatterplot smoothing (LOWESS) line is shown in red (frac=0.5). Absolute Spearman’s rank correlation coefficient (ρ) is shown for each plot; for *KCNQ4*, the correlation was estimated on absolute deviation from typical values. **C)** Plots in ‘B’ are repeated using REVEL scores instead of AlphaMissense scores. **D)** The relationship between AlphaMissense and REVEL scores is shown for each gene.

Variants with low AlphaMissense scores are generally found to be non-functional (Fig. 2B) and overlap with “Benign” and “Likely Benign” ClinVar labels (Fig. 2A). However, performance is worse for variants with high AlphaMissense scores. Many variants did not impact function in these assays (Fig. 2B) and were not labeled “Pathogenic” or “Likely Pathogenic” in ClinVar (Fig. 2A), suggesting overcalling of pathogenic variants. This overcalling issue is not limited to AlphaMissense, being also observed in REVEL (Fig. 2C) and multiple other missense severity metrics (Table S1).

## Discussion

Reliable prediction of missense variants is a complex problem that remains incompletely solved. AlphaMissense demonstrates that there is still scope for computational approaches leveraging orthogonal data, such as protein structure, to improve on existing algorithms. Based on an analysis of five genes, we find AlphaMissense to have the best correlation with functional data across the 17,696 variants, though the improvement over other algorithms is modest (Fig. 1, Table 1). Compared with functional data, AlphaMissense and many other algorithms tend to overcall pathogenic variants (Fig. 2B/C). This may reflect a limitation of the functional assays (e.g., the missense variant impacts an alternative role of the protein or a cell-type specific effect); however, the ClinVar data are generally supportive of the functional data (Fig. 2A).

Predicting missense variant pathogenicity is further complicated by both how the missense variant impacts protein function and the effect size of disruption of the gene on a phenotype in a population (e.g., penetrance). Metrics driven by selective pressure, such as conservation across species or genomic constraint, tend to integrate the combination of protein function and gene disruption effect size. In contrast, measures of protein stability or structure alone might be highly predictive of protein function independent of gene disruption effect size. Mode of inheritance will further modulate these considerations. Additionally, no proteome-wide algorithms are capable of reliably distinguishing loss-of-function from gain-of-function variants, which may be critical for precision therapeutics.

At present, the major limitation in algorithm design is likely to come from the paucity of training data, with only a tiny fraction of possible missense variants having been characterized reliably. Increasing the size and ancestral diversity of human population cohorts will help improve these algorithms, but there may be no alternative to performing MAVE on many more genes.^17^

## Supporting information

Missense_variants

## Acknowledgements

This work was supported by grants from the National Institute of Mental Health (R01MH125516, R01MH122681 and R01MH129751 to SJS).

## Declaration of interests

Stephan Sanders receives research funding from BioMarin Pharmaceutical.

